# Microbiota Dynamics in Lionfish (Pterois): Insights into Invasion and Establishment in the Mediterranean Sea

**DOI:** 10.1101/2025.02.03.635474

**Authors:** Dalit Meron, Maya Lalzar, Shevy Bat-Sheva Rothman, Yael Kroin, Elizabeth Kaufman, Kimani Kitson-Walters, Tal Zvi-Kedem, Eli Shemesh, Rami Tsadok, Hagai Nativ, Shai Einbinder, Dan Tchernov

## Abstract

Lionfishes (*Pterois* spp.), originally native to the Indo-Pacific and Red Sea, have become one of the most invasive marine species globally, including the recent establishment in the Mediterranean Sea. This study investigates the microbiota of lionfish to explore its potential role in their invasion success and establishment. Using high-throughput sequencing and microbiota analyses, we characterized the species-specific core microbiome and identified habitat-specific markers across different regions (Red Sea, Mediterranean Sea, Caribbean, and aquarium populations) and organs. Focusing on the Mediterranean invasion, we tracked lionfish distribution and population dynamics along the Israeli coastline from 2017 to 2023, monitoring size, seasonal trends, and depth preferences. Our findings reveal that lionfish initially established themselves in deeper waters before expanding to shallower habitats, with a gradual increase in population size and body length over time. From a microbial aspect, we compared the microbiota of lionfish organs and identified a similar pattern (*Photobacterium)*, to Earlier Lessepsian migrants fish species. This study provides novel insights into the interactions between microbiota and host ecology, shedding light on the mechanisms that may support the successful invasion. This study contributes to the understanding of lionfish invasion dynamics in the Mediterranean. It highlights the microbiota as an integral component for studying the ecological and biological mechanisms underpinning invasive species’ success and establishment of lionfish.

## Introduction

Lionfish (LF) species (*Pterois volitans* and *P. miles*), are reef fish native to the Indo-Pacific and Red Sea, are considered among marine ecosystems’ most notorious invasive species (Côté & Smith 2018). They have become significant and successful invaders with global distribution, including the northwestern Atlantic, the eastern coast of the USA, Bermuda, the Caribbean region and the Gulf of Mexico (Schofield, 2009; Côté & Smith, 2018), Brazil (Ferreira et al., 2015) and more recently, the Mediterranean Sea (Kleitou et al., 2016). In the Mediterranean Sea, *P. miles* were first recorded near Israel in 1991 (Golani & Sonin, 1992), and the invasion notably accelerated from the mid-2010s and onwards. The Suez Canal has served as a key pathway for this introduction, also termed “Lessepsian migration” facilitating the species’ establishment in new habitats (Bariche et al., 2017; Stern et al., 2019). The unprecedented speed and scope of their spread highlight their adaptability and the significant impact they impose on invaded ecosystems.

Several biological and ecological factors contribute to the LF’s invasion success (Côté & Smith 2018; Bottacin et al., 2024). Lionfish are generalist predators whose diet includes a wide variety of fish and invertebrates. A stomach content analysis has revealed that LF in the Mexican Caribbean preyed on nearly 50 different local fish species (Arredondo-Chávez et al., 2016), underscoring their lack of dietary selectivity and facilitating their adaptation to new environments. Their early maturation (within the first year) combined with a robust strategy (asynchronous egg release) and fast growth, contribute to the rapid increase in LF density and their successful establishment in invaded areas (Morris et al., 2011; Côté et al., 2013; Albin 2013). Additionally, the absence of natural predators in the invaded regions allows LF populations to grow unhindered, thus significantly disrupting the local marine ecosystems. This invasion leads to a decline in local species and changes biodiversity and community structure, affecting the reef ecosystems’ resilience (Albins & Hixon, 2008; Green et al., 2012; Côté et al., 2018).

Invasive species are often characterized by their remarkable adaptability, which enables them to thrive in novel environments with different abiotic and biotic conditions compared to their native habitats. The microbiota may influence various host functions, including immune responses, nutrient acquisition, and environmental resilience, providing an invasive species with significant advantages in unfamiliar territories (Aires et al., 2016; Ronanki, 2024; Coats et al., 2014). Another aspect concerns the symbionts of invasive species in the new environments. Numerous studies have shown that various stressors, such as temperature changes, pollution, and fluctuations in resource availability, can potentially impact the dynamics of bacterial invasion in aquatic ecosystems (Liu et al., 2012; Bagra et al., 2022). These stressors may not only affect the composition of bacteria and disrupt established symbiotic relationships but also create new niches that invading microorganisms can exploit (Amalfitano et al., 2015; Cavicchioli et al., 2019; Paerl et al.,2016).

Although research has explored the effects of environmental changes on microbiota, the specific role of microbiota in supporting the survival and establishment of invasive species remains inconclusive. The role of symbiotic bacteria in invasive species has been investigated mainly in terrestrial ecosystems (Lu *et al*., 2016; Bang *et al*., 2018). Research focused on insects for example, has demonstrated that altering the microbiota can influence various traits, including: suppressing native competitors (Vilcinskas et al., 2013), enhancing nutrient acquisition (Adams et al. 2011), promoting pest resilience (Brown et al., 2014) and increasing fecundity and survival rates (Himler et al., 2011). These traits contribute to establishing and spreading invasive species within new environments. However, only a few studies have demonstrated the contribution of the microbiota to invasion within marine ecosystems. Among them, a study on the invading alga *Caulerpa racemosa* showed that an increase in bacteria involved in sediment biogeochemical processes contributed to the invasion’s success by displacing native seagrasses (Aires et al., 2013). Another study demonstrated that symbiotic relationships between chemoautotrophic bacteria and marine invertebrates can influence metabolic capabilities and expand the adaptation to a variety of ecological niches for the host (Lee et al., 1999). Thus far, only a few studies have examined the LF microbiota aspect and have mainly focused on the Indo-Pacific (native) and western Atlantic (invaded) regions (Stevens & Olson, 2013; Stevens & Olson, 2015; Stevens et al., 2016). However, these were conducted about a decade ago, using methods that were less advanced and comprehensive than those available today, such as high-throughput amplicon sequencing and metagenomics

Invasive species may serve as ideal models for investigating the host-microbe-environment interactions. By comparing the microbiota in native and invaded environments, we can investigate three aspects: 1) how the new environmental conditions shape and change the composition and diversity of the bacterial communities; 2) what is the stable core microbiota (that persists despite the changing conditions), and 3) to examine whether the microbiota impacts the success of invasion. This study compares the microbiome aspect of LF from different origins: The Mediterranean and the Caribbean (invaders), the Red Sea (native), and local aquarium. In addition, we chose to focus on the LF in the Mediterranean Sea, a relatively recent and successful invader, making it an ideal case study for examining its invasion, establishment, and interactions with microbial symbionts. To this end, fish surveys, which included fish counts and visually estimated lengths, were conducted over a five-year period. In parallel, for comparison, microbial profiling and trophic position (TP) of the two earlier Lessepsian invasive species, *Sargocentron rubrum* and *Siganus rivulatus*, were also described.

## Methods

### Fish sampling

Between 2019 and 2022, a total of 90 fish specimens were sampled, comprising five individuals of *S. rubrum*, nine of *S. rivulatus*, and 79 of *Pterois* spp. (lionfish) (Table S1). Lionfish were collected from the Gulf of Eilat in the Red Sea (RS), the Mediterranean coast of Israel (MS), along the Israeli coast, Sint Eustatius in the Caribbean Sea (CA) and from captive populations in the Israel Aquarium in Jerusalem (AQ_J) (https://www.israel-aquarium.org.il). LF from the latter were originally captured off the coast of Kenya and kept in the aquarium since 2018. Fish specimens from the wild (from MS, RS and CA) were caught using spearguns and transported on ice to the laboratory. In contrast, captive fish specimens were collected with (hand) nets, sampled immediately, and returned to the aquarium. Size (total length) and sex (when applicable) indices were recorded for each individual. Sterile swabs were used to collect samples from four organs of each specimen: cloaca, mouth, gills, and skin. Surrounding water was also sampled, 1.5 liters filtered using a single-use “Nalgene RapidFlow Filters” 0.2μm (Thermo Scientific, cat no. 566–0020, Israel). The swabs and filters were stored at −20°C for further analysis. In total, 378 (365 fish and 13 water) samples were collected and analyzed (Table S2).

### Lionfish surveys in the Mediterranean Sea

As part of the Morris Kahn Marine Research Station (MKMRS) monitoring program, fish surveys were conducted by SCUBA in central (Sdot-Yam) and northern (Achziv) Israel during the Spring (April to May) and Fall (October to November) at depths of 10, 25, and 45 meters. At each depth, an area of 250 square meters was surveyed. Surveys included fish counts and visually estimated lengths. The data on LF from these surveys were analyzed for the period of 2017 - 2023. The fish survey method and data are available at https://med-lter.haifa.ac.il/database-hub/.

### DNA extraction, PCR amplification and amplicon sequencing

DNA was extracted from all samples using the DNeasy PowerSoil Pro Kit (Qiagen, www.qiagen.com) following the manufacturer’s instructions. Partial sequences of the 16S rRNA gene at the V4 hypervariable region were amplified using the primers 518F (ACCAGCAGCCGCGGTAATACG) and 806R (GGACTACNVGGGTWTCTAAT) (based on Apprill *et al*.., 2015; Parada *et al.,* 2016 with few modifications) that contained 5’ common sequence tags (CS1 and CS2) (Moonsamy et al., 2013). Amplicons were generated using a two-stage PCR amplification protocol described by Naqib et al. (2018). Cycling conditions for the first stage PCR were 94°C for 15s, 50°C for 20s, and 72°C for 20s for 15 cycles, with an additional 15 cycles with 62°C for 15s (annealing stage). Subsequent steps were carried out at the Genome Research Core (GRC) within the Research Resources Center (RRC) at the University of Illinois at Chicago (UIC). A second amplification was performed for each sample, with a separate primer pair with a unique 10-base barcode obtained from the Access Array Barcode Library for Illumina (Fluidigm, South San Francisco, CA; Item# 100-4876). Cycling conditions were 95 °C for 5 minutes, followed by eight cycles of 95 °C for 30 sec, 60 °C for 30 sec and 72 °C for 30 sec. The amplified barcoded PCR products were pooled and purified using an AMPure XP cleanup protocol (0.6X, vol/vol; Agencourt, Beckmann-Coulter). With a 15% phiX spike-in, the pooled libraries were loaded onto an Illumina MiniSeq mid-output flow cell and sequence (2×153 bases paired-end reads). Barcode sequences were used for sequence read de-multiplexing of raw data, which was then recovered as FASTQ-formatted files. Raw sequence data is available in the NCBI SRA database under PRJNA1213691.

### Sequence processing

The Dada2 pipeline (https://www.nature.com/articles/nmeth.3869) (dada2 package version 1.20.0) was used for sequence data processing. The processing was conducted for each set of samples and each run separately for the following steps: Sequences were filtered and trimmed for quality using the ‘filterAndTrim’ command with the parameters maxN set to zero, maxEE set to 2, trimLeft set to 21 and 20 bases for the forward and reverse reads, respectively. The sequence error estimation model was calculated using the ‘learnErrors’ option, and then, the dada2 algorithm for error correction was applied using default parameters. Forward and reverse reads were then merged with minimum overlap set at eight bp. Suspected chimaera was detected and removed using the command ‘removeBimeraDenovo’, and a count table was produced. To obtain a taxonomic assignment for each ASV, ASV sequences were aligned to the ARB-Silva small subunit rRNA database (version Silva_nr_138.1) using the command ‘assignTaxonomy’ with default parameters. However, minBoot set at 80% (Table S2).

### Data analysis

Microbiota analysis procedures were conducted in R (version 4.3.1). For analysis of microbiota composition, counts data had been normalized by the cumulative sums squares (CSS) method using the R package ‘metagenomSeq’ (version 1.42.0). Normalized counts were then summed sample-wise, based on taxonomic assignment, up to the genus taxonomy level. Core microbiota composition was determined based on the prevalence of the different genera detected. A genus was considered a core member if its prevalence at each site tested was above 25%. To test the effect of the collection site (Sea) or organ on the composition of the microbiota, a permutational analysis of variance (PERMANOVA) test was performed using the ‘adonis2’ function in the R package ‘vegan’ (version 2.6.4). Non-metric multidimensional scaling (NMDS) ordination was calculated using the R package ‘vegan’ based on Bray-Curtis dissimilarities. To extract microbiota markers, Linear Discriminant Analysis (LDA) Effect Size (LEfSe) was calculated using the function ‘run_lefse’ from package ‘microbiomeMarker’ (version 1.6.0). The association was significant for Bejnamini-Hochberg adjusted *P* value <0.05 and LDA effect size of 2. For the calculation of the effect of sex, PERMANOVA was performed separately, using 112 samples corresponding to 34 fish specimens for which sex could be determined. Additionally, the contribution of sex to variance was tested by canonical correspondence analysis (CCA) in ‘vegan’ considering the model microbiota∼sex+site+organ. CCA model and items were considered significant at *P*<0.05.

For alpha-diversity parameters calculations, the count data was subsampled to 4,000 reads per sample using the function ‘rrarefy’ from the R package ‘vegan’ package (version 2.6.4). Using the rarified data, Shannon H’s index of diversity, and the Simpson’s index for evenness and richness, calculated as observe number of genera were calculated with ‘vegan’. To test the effect of site, organ and their interaction on alpha diversity parameters, the aligned rank-transformed (ART) ANOVA test was applied, with the R package ‘ARTool’ (version 0.11.1). Post-hoc pair-wise tests were also conducted with package ‘ARTool’ using the ‘art.con’ function and the Benjamini-Hochberg method for *P* value adjustment against false discovery.

## Results

### Lionfish microbiota composition

Lionfish microbiota composition was compared among four sites (MS, RS, CA and AQ_J) and among the four organs: mouth, gills, cloaca, and skin (Total 307 samples; Table S1). To define the LF core microbiota, we examined the prevalence of the different genera identified and considered those with prevalence > 25% at all of the examined sites. Based on these criteria, 18 bacterial genera were recognized as the LF core microbiota (Fig. 1A, Table S2). The most prevalent genera were *Photobacterium*, *Vibrio*, *Pseudomonas* and *Shewanella* (prevalence above 80% of the samples), all belonging to the class Gammaproteobacteria. While a robust core microbiota was thus suggested, an inspection of the dominant genera in each organ at each site (> 4% mean relative abundance) indicated major differences in composition between sampling sites (Fig. 1B). The contribution of the site to variation in microbiota composition, as assessed by PERMANOVA was highly significant (*P* < 0.001, R^2^=0.133). Pair-wise comparisons further indicated unique microbiota composition for each site (Table S3). This was corroborated by NMDS ordination (Fig. 2A). Linear decomposition model (LDA) effect size analysis was used to identify key genera as markers of each site (Table S4_LDA). The most noted genera identified (LDA score >3) were *Psyhromonas* (AQ_J), *Pseudomonas* (CA), *Photobacterium* and *Vibrio* (MS), *Cetobacterium*, including *Clostridiaceae* and *Caedibacter* (RS) (Fig. 2B). The contribution of the organ to variation was examined for each site separately, using PERMANOVA. Organ effect was significant for LF sampled at site RS (*P* <0.001, R^2^=0.087) and CA (*P* <0.001, R^2^=0.066), as was indicated from inspection of composition (Fig. 1B). Post-hoc pairwise tests for organs in RS concluded a significant difference between each pair, with the greatest effect between the gills and the cloaca microbiota (gills-cloaca: *P*< 0.001, R^2^=0.089; gills-mouth: *P*=0.005, R^2^=0.041; gills-skin: *P*< 0.001, R^2^=0.072; mouth-cloaca: *P*<0.001, R^2^=0.066; mouth-skin: *P*=0.006, R^2^=0.046; skin-cloaca: *P=*0.022, R^2^=0.039) while in site CA significant difference was observed only between skin and cloaca (*P* <0.029, R^2^=0.039).

**Figure 1:**
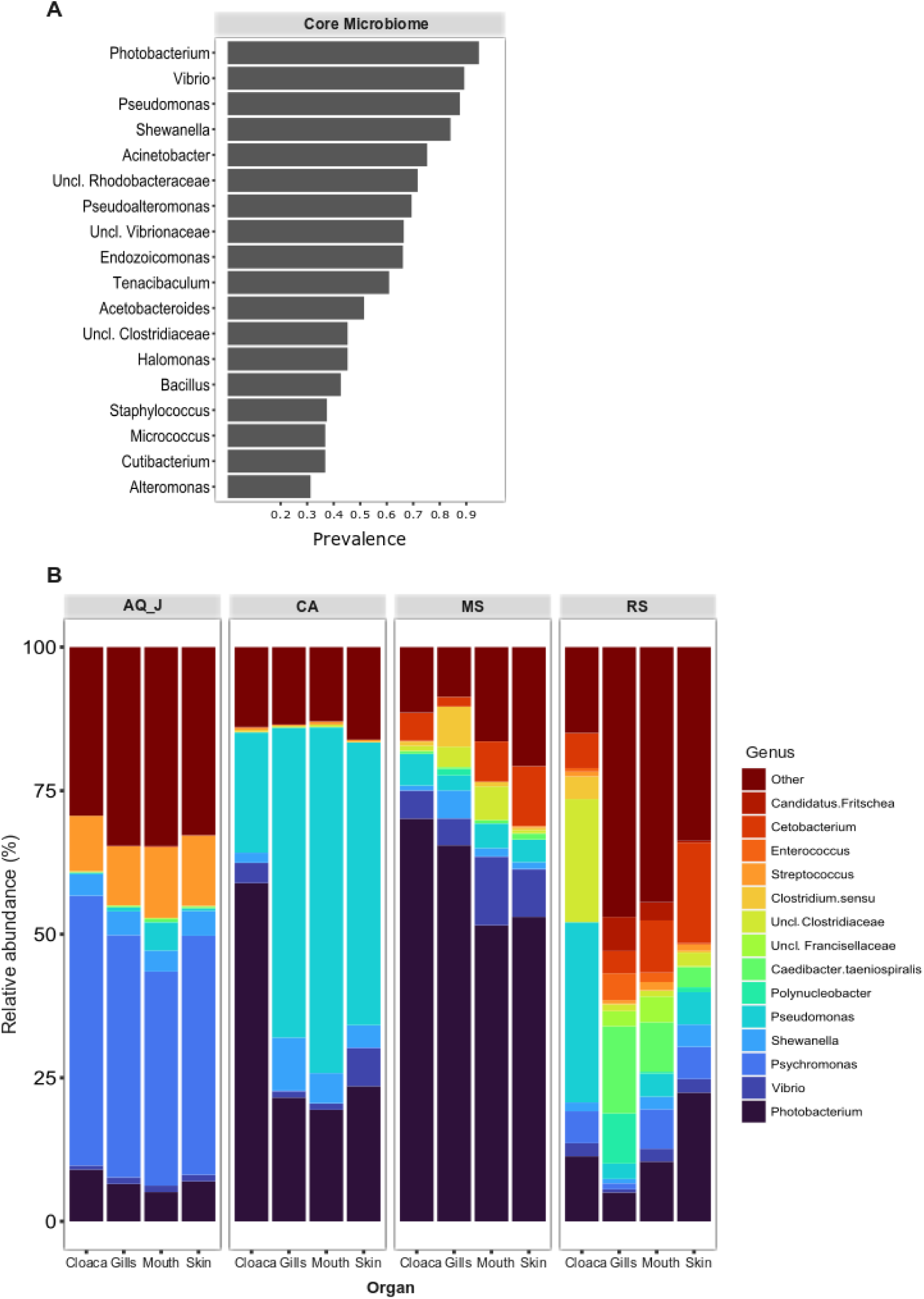
Composition of lionfish microbiota. Lionfish microbiota was determined by amplicon sequencing of 16S rRNA gene fragments from total DNA extracted from 307 samples, representing the gills, cloaca, skin and mouth of specimens collected at the Mediterranean Sea (MS), Red Sea (RS), Caribbean Sea (CA) and the Jerusalem aquarium (AQ_J). (**A)** Core microbiota composition. Includes genera for which prevalence was >25% at each of the sampled sites. (**B)** Composition of dominant genera (relative abundance >5% at least at one site).

**Figure 2:**
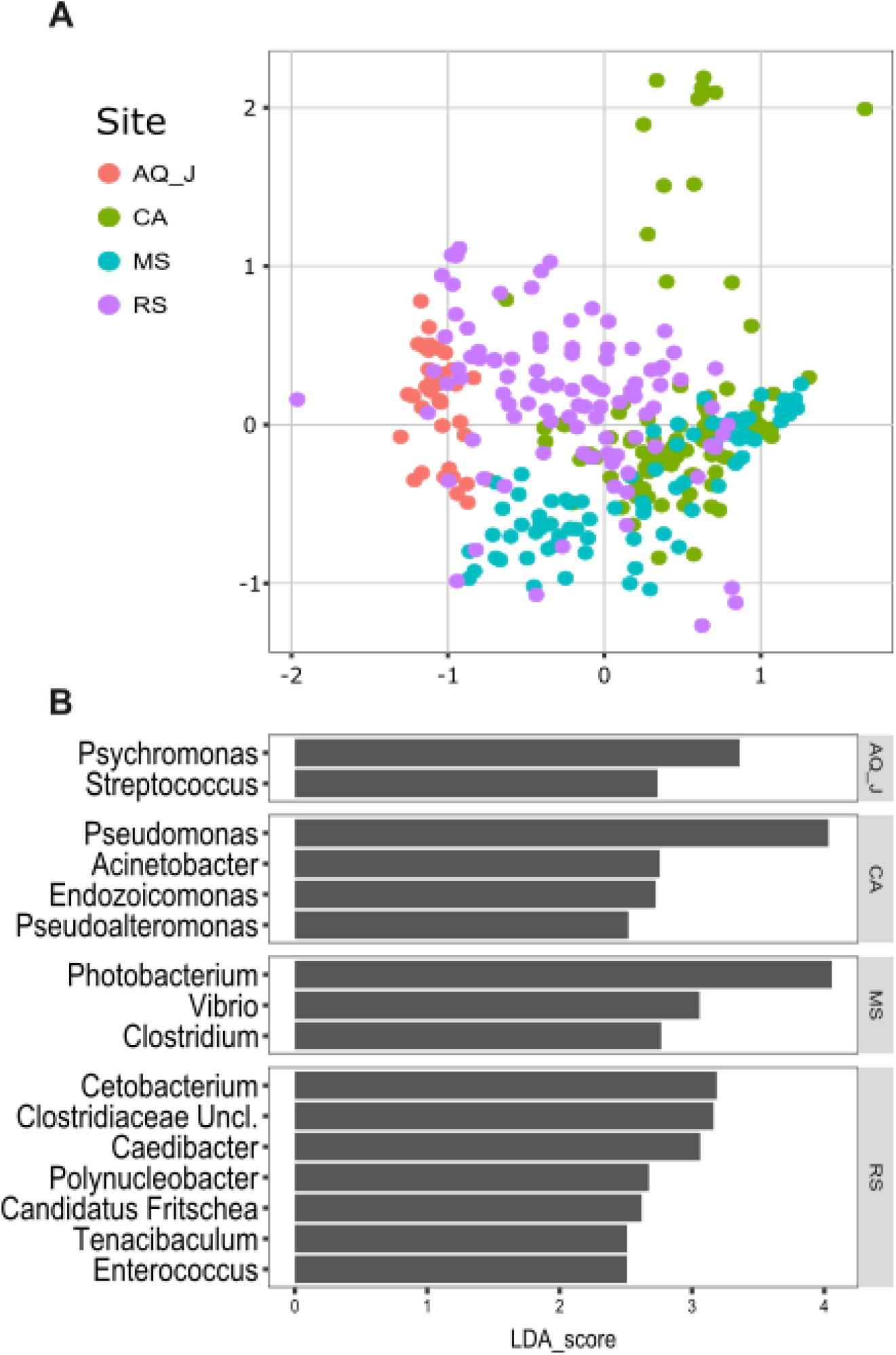
Variation in lionfish microbiota composition among sites. (**A)** Non-metric multidimensional scaling (NMDS) ordination, based on Bray-Curtis distances (Stress*_K=3_*=0.16). (**B)** Microbial markers for sites, based on linear decomposition model effect size (LEfSe) analysis. Presented are significant markers (Benjamini-Horchberg adjusted P value <0.05) for which the effect size was >2.5.

The effects of site and organ were also reflected in microbiota diversity, examined at the genus level of taxonomy. ART-ANOVA test confirmed the effect of the site, without significant interaction for Shannon H’ and Simpson indices (Table 1).

**Table 1:**
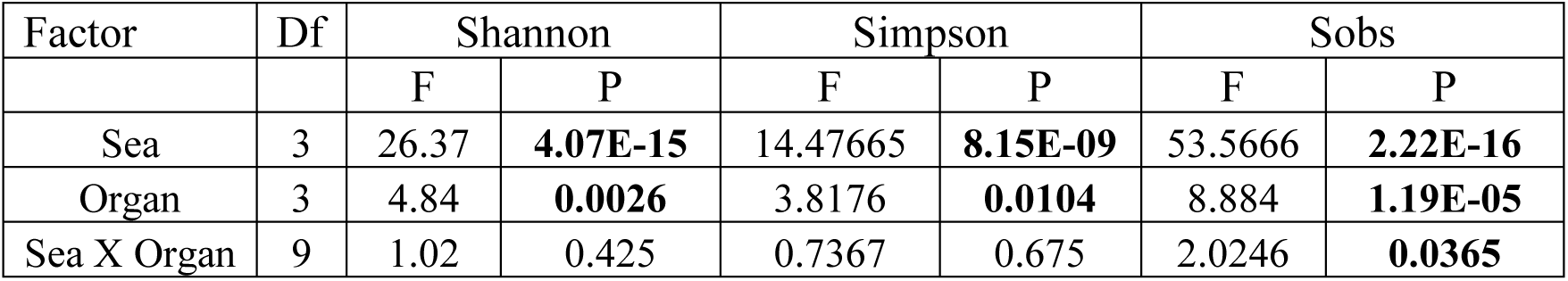
Effect of site and organ on lionfish microbial community structure. Alpha diversity indices Shannon H’, Simpson and richness (the number of species observed (Sobs)) were calculated and compared by aligned rank transformed ANOVA.

| Factor | Df | Shannon |  | Simpson |  | Sobs |  |
| --- | --- | --- | --- | --- | --- | --- | --- |
|  |  | F | P | F | P | F | P |
| Sea | 3 | 26.37 | **4.07E-15** | 14.47665 | **8.15E-09** | 53.5666 | **2.22E-16** |
| Organ | 3 | 4.84 | **0.0026** | 3.8176 | **0.0104** | 8.884 | **1.19E-05** |
| Sea X Organ | 9 | 1.02 | 0.425 | 0.7367 | 0.675 | 2.0246 | **0.0365** |

Richness, calculated as the observed number of genera, was significant for site and organ as well as their interaction Based on pairwise comparisons among sites the CA site had the lower diversity compared to all other sites, and MS diversity was lower than that of AQ_J. Pairwise comparison among organs calculated significantly lower diversity for the cloaca compared to the mouth and the skin (Table S5). Another aspect examined for effect on microbiota composition was sex. The sex could be identified for a subset of 34 fish specimens (112 samples). PERMANOVA test indicated no significant effect for sex. However, canonical correspondence analysis (CCA) did significantly link sex to microbiota composition (χ^2^=0.31, *P*=0.005) and explains ∼1.1% of inertia (Fig. 3).

**Figure 3:**
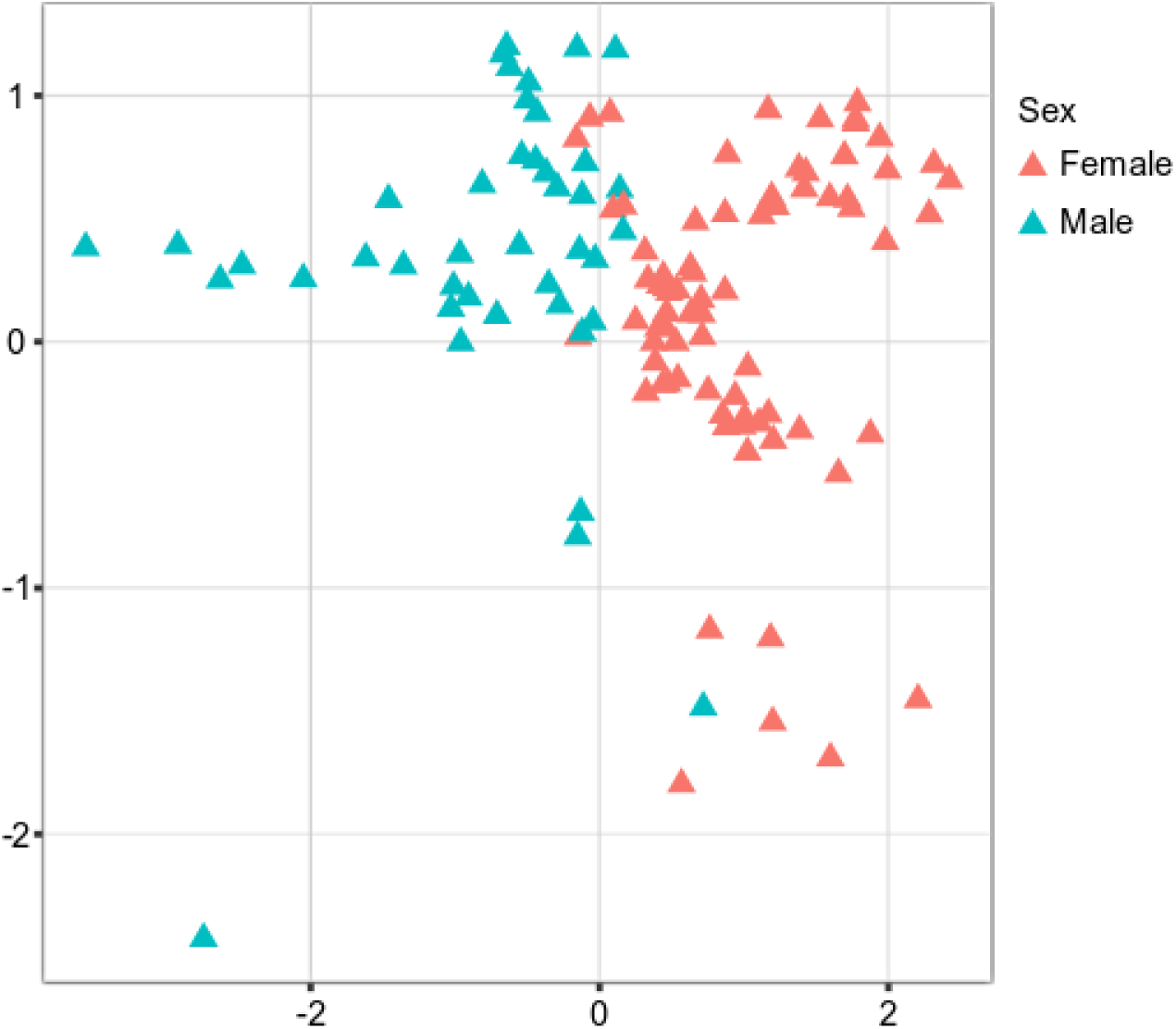
Contribution of sex to variation in lionfish microbiota composition. Canonical correspondence analysis (CCA) analysis determined sex as a significant factor (*P*<0.05) contributing 2.8% of variance in microbiota composition. Presented is CCA ordination plot.

### Lessepsian invasive fish (RS - MS)

Interestingly, when analyzing the bacterial overlap at the genus level across paired organs, it was found that over 67% of the bacteria identified in MS were shared between the two regions. Conversely, a notably higher percentage of bacteria was unique to RS, with 47-41%, compared to only 22-23% of unique bacteria in MS (Fig. 4). Noting the major difference in the relative abundance of the *Photobacterium* in the MS vs. RS samples (Figs. 1B and 2B), we extended our microbiota comparison to additional Lessepsian invasive fish species. We chose *Sargocentron rubrum* and *Siganus rivulatus*, known as Earlier Lessepsian migrants (*S. rubrum* reported by Haas & Steinitz in 1947 and *S. rivulatus* by W. Steinitz in 1927), and characterized their microbiota. Overall relative abundance of *Photobacterium* ranged differently levels among species (0.6% - 59.2% for *S. rubrum*; 0% - 24.2% for *S. rivulatus*). Nevertheless, regardless of the organ, *Photobacterium* was much more dominant in MS specimens compared to RS ones (Fig. 5). Interestingly, the *Photobacterium* relative abundance was negligible (less than 1%) in the MS water samples (Table S2).

**Figure 4:**
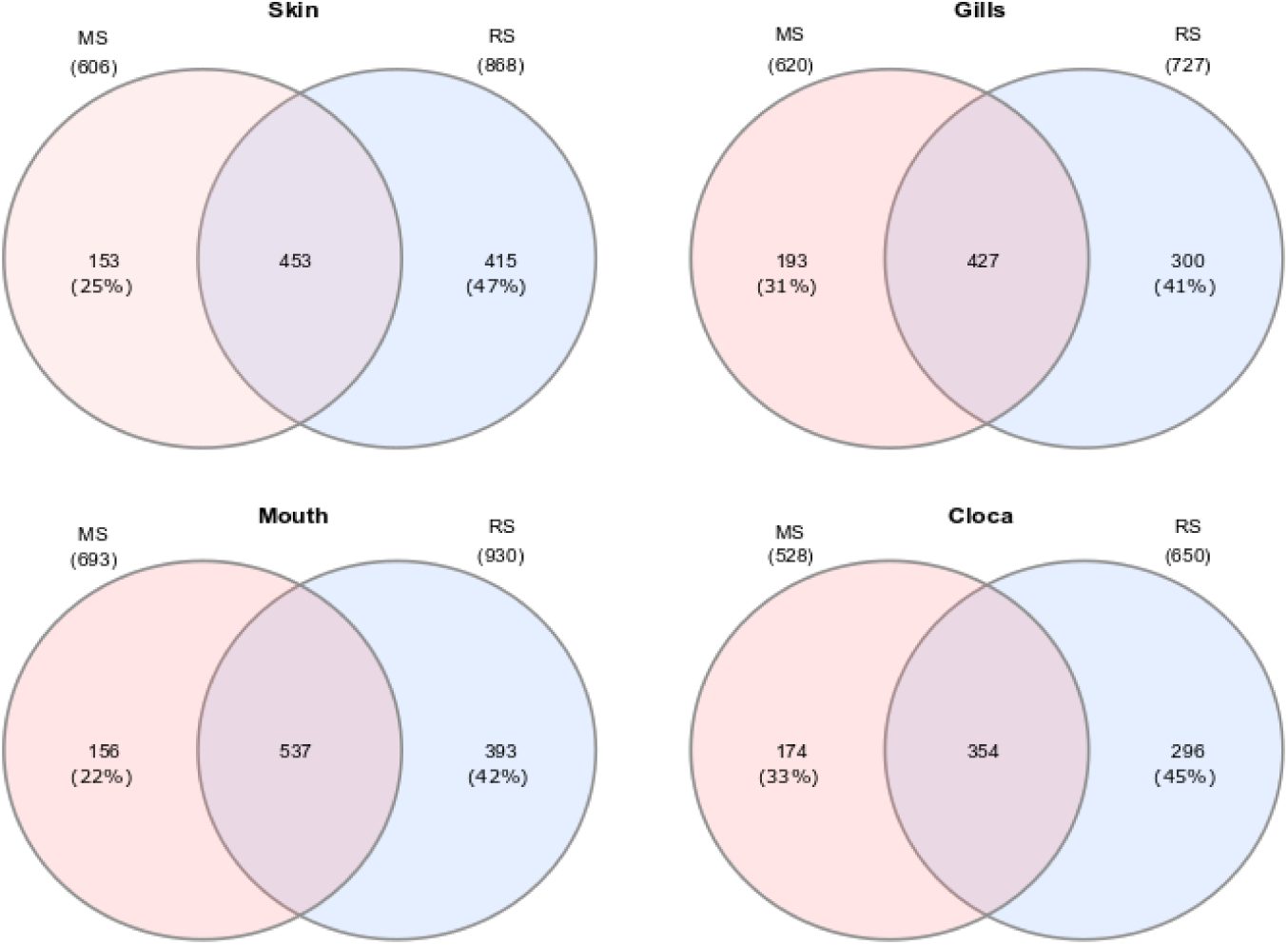
Venn diagrams presenting shared and unique microbial genera among sites for different organs of Lionfish. Numbers in brackets represent the percentage from total number identified genera at the specific site and organ.

**Figure 5:**
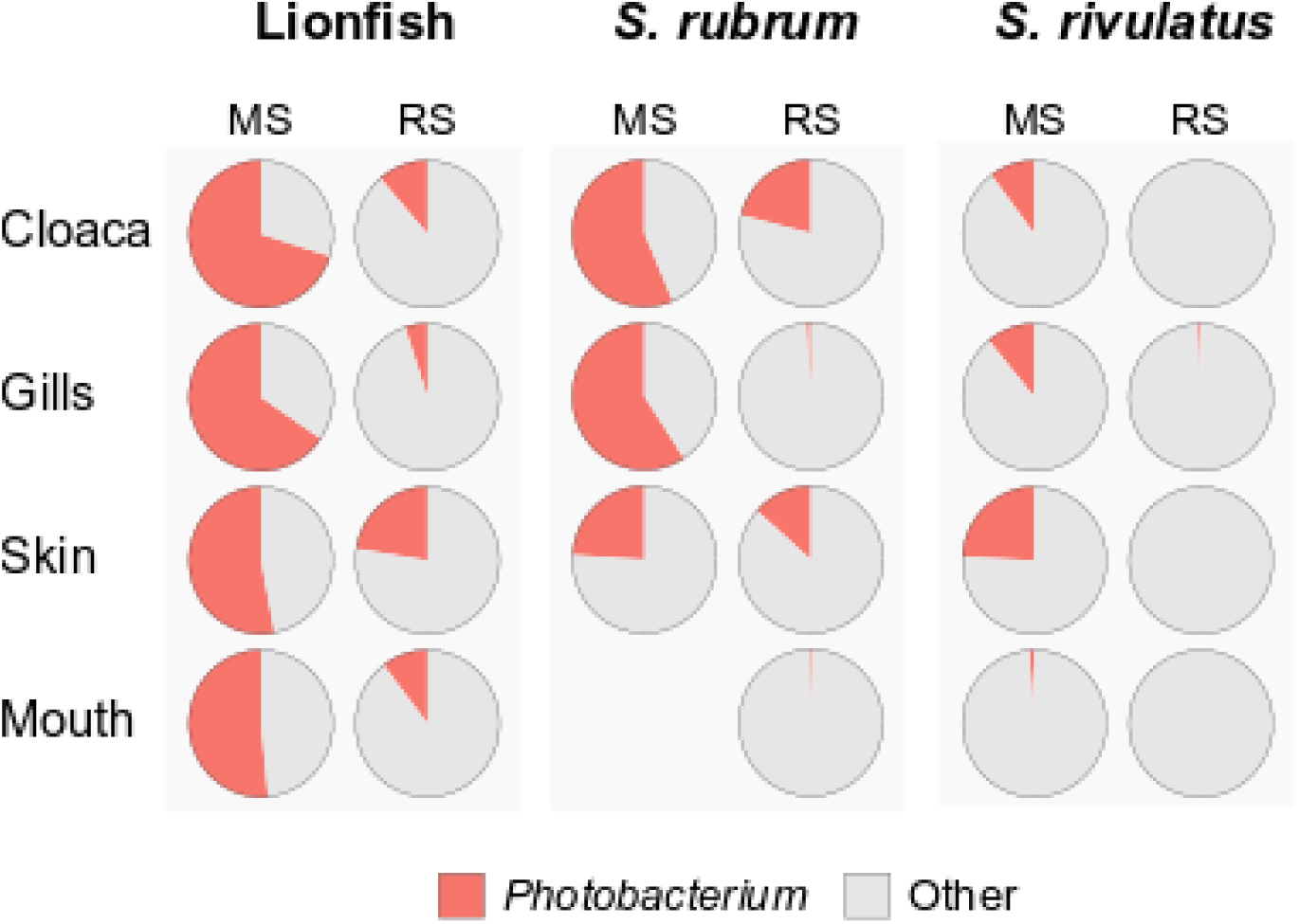
Dominance of *Photobacterium* across species, sites and organs. Relative abundances of *Photobacterium* are presented as pie charts.

### Lionfish MS - invasion and establishment

Based on the fish surveys, we analyzed the LF data and compared the number of fish observation and their size by site, year, season, and depth (Fig. 6). LF were first recorded in fish surveys in 2017 in the northern site (Achziv). The first sightings of LF in the Sdot-Yam area occurred only two years later. At both sites, initial observations of LF were recorded from the deepest transect (45 m), with sightings at shallower depths (25 m and 10 m) appearing only two to three years later. A consistent upward trend in LF population size was observed at Achziv. In contrast, no such trend was noted at Sdot-yam (Fig. 6A). Nevertheless, at both sites, the majority of observations were in the fall (Fig. 6B).

**Figure 6:**
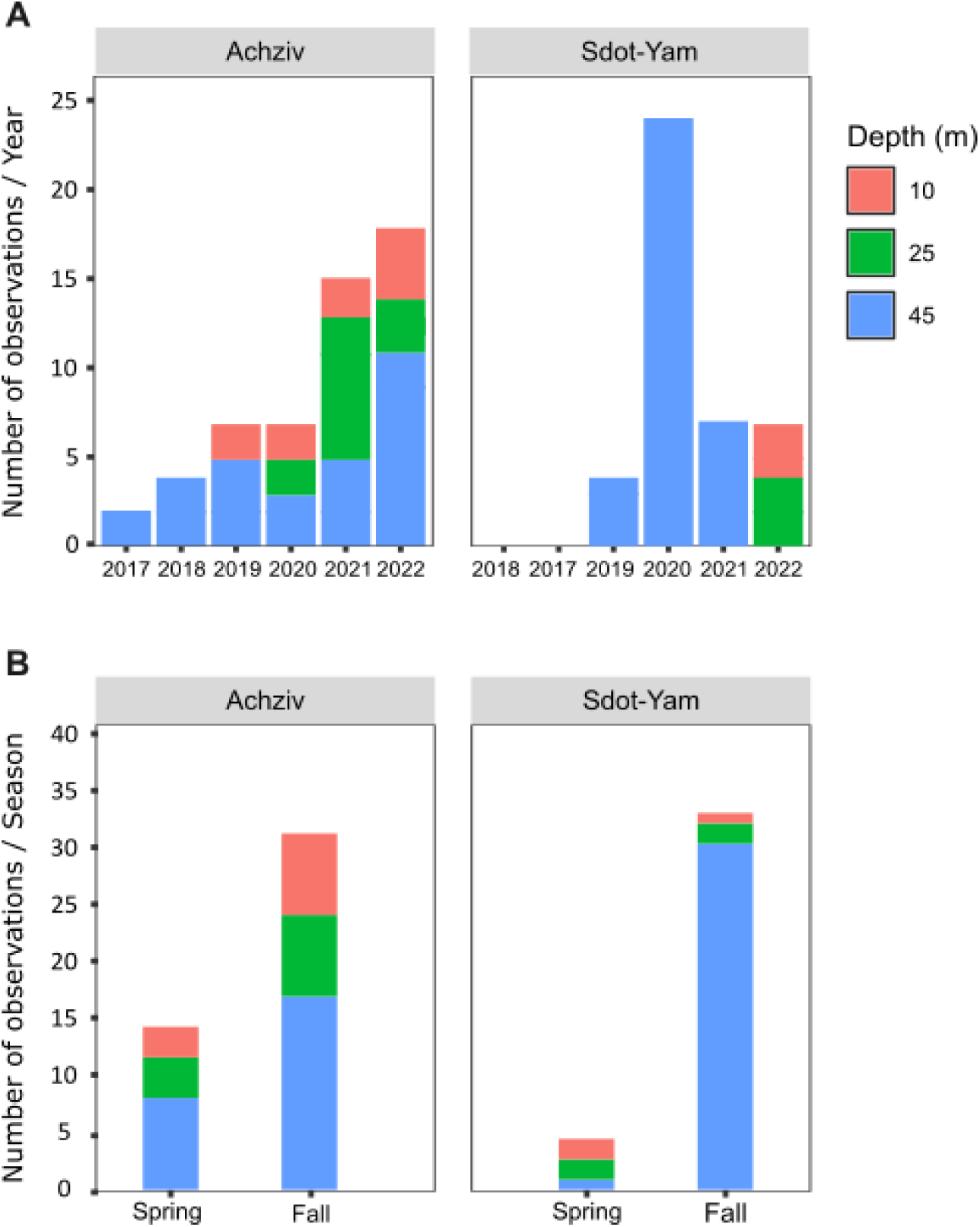
Lionfish establishment in the Mediterranean Sea (2017-2022). Numbers of observations of lionfish at different seafloor depths along the Achziv and Sdot-Yam transects **(A)** or at different seasons **(B)**. Taken from MKMRS database https://med-lter.haifa.ac.il/database-hub/

Fish surveys also assessed the approximate size of the fish (in terms of length) and indicated a substantial increase in size observation between 2017 and 2023 (Fig. 7A). Length measurement of collected fish specimens from MS (2019-2021), RS (2020- 2022) and CA (2021) confirmed that during the first years of invasion, LF at MS were smaller in size compared to RS or CA (Fig 7B). Considering that smaller sizes may be accounted for by the quality of nutrition, we examined the TP value. In LF, the value of TP was the lowest in the CA. However, no differences were observed between RS and MS (Fig. 7C). Interestingly, the opposite trend was found for the two other Lassepsian invaders examined, *S. rubrum* and *S. rivulatus,* TP values were lower in MS compared to RS.

**Figure 7:**
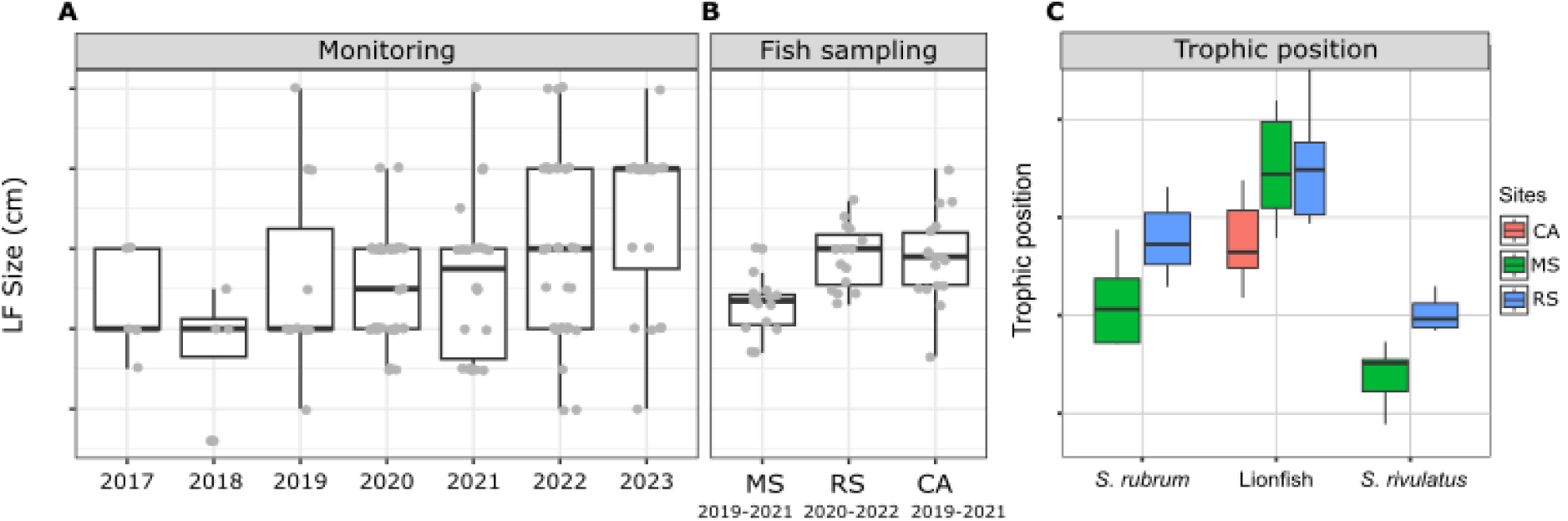
Lionfish establishment at the Mediterranean Sea (2017-2023). **(A)** Size estimate distribution from visual fish surveys. **(B)** Size distribution of captured lionfish specimens at different sites. **(C)** Trophic position of captured lionfish specimens, calculated based on Trophic isotopic discrimination of amino acids (glutamic acid and phenylalanine) (Takizawa & Chikaraishi, 2024).

## Discussion

Lionfish are widely regarded as one of the most invasive and successfully established species in the marine environment. Originally native to the Indo-Pacific Ocean and the Red Sea, they have spread across the entire tropical western Atlantic, including the Brazilian coast and the Mediterranean Sea (Côté & Smith 2018; Soares et al., 2022, 2023; Kletou et al., 2016). As previously described, various parameters such as the ability to survive, reproduction rate, the absence of predators, and its adaptation to a wide range of conditions, contributed to the success of its invasion and establishment (Bottacini et al., 2024). In this study, we focused on the microbiome aspect of LF, aiming to characterize the core microbiome shared by the species and to examine differences and variations across different habitats. We also followed the process of the Lessepsian migration and the establishment of LF populations in the Mediterranean Sea.

### Lionfish microbiota

The microbiota of invasive species is crucial to their success and interactions in new ecosystems. Microbial diversity and functional stability can influence how likely an invasive species is to establish itself (Amalfitano, 2015; Huang 2019). We characterized the core microbiome of LF based on samples collected from four sites, and various organs. The Gammaproteobacteria class exhibited the highest relative abundance within the core microbiome, with dominant genera such as *Photobacterium*, *Vibrio*, *Pseudomonas*, and *Shewanella* (Fig. 1A and 2C). The information on the microbiota of LF is very limited; however, Stevens and Olson (2015) and Stevens et al. (2016) reported similar findings regarding the bacterial groups in LF from the Indo-Pacific and Western Atlantic. Notably, *Pseudoalteromonas* and *Photobacterium*, identified as key markers in the invaded regions (CA and MS, respectively, Fig. 1B) are known to exhibit broad antibacterial activity. Compared to that of local fish, this robust antibacterial capability may provide LF with a competitive advantage, enhancing their resilience and success in new habitats (Stevens et al. 2016).

Various environmental parameters, such as temperature, pH, and salinity, have been shown to significantly influence the diversity and abundance of microbiota (Thompson et al., 2017; Coutinho et al., 2015; Hicks et al., 2018). In this study, we observed notable differences in microbiota composition across the analyzed sites, with each site characterized by unique bacterial markers. Identifying site-specific markers may facilitate tracing the origin of LF habitats, providing a valuable tool for monitoring their distribution. This knowledge is particularly relevant in cases of species invasion, where understanding habitat origins can inform management strategies and control measures. In the RS, which represents a native habitat for LF, we identified the highest number of significant microbial markers (with scores exceeding 2.5). This high number of markers may also reflect the influence of the organ factor, which contributed to the microbiota variation in this region, adaptation and fostering niche-specific microbiota associations within different organs (Kuang et al., 2020; Meron et al., 2020). In contrast, fish introduced to new habitats, such as in invasive contexts, may only develop distinct organ-specific microbiota at later stages of establishment. The unique and diverse bacterial communities in each organ of RS lionfish suggest the evolution of stable and well-adapted microbiota, shaped over generations by local environmental pressures. In the CA region, a partial trend was observed, potentially reflecting an intermediate stage in establishing lionfish populations. First documented in the Caribbean in the early 2000s, lionfish were officially recognized as a successful invasive species around 2004– 2005 (Schofield, 2009). This indicates that LF in the CA region are beginning to adapt to the local environment. By contrast, the relatively recent invasion of LF in the MS region suggests that adaptation and microbiome stabilization are still in their early stages.

In the case of LF from AQ_J, although their origin traces back to the coast of Kenya (a native habitat for LF), the aquarium environment—characterized by artificial water in a closed system—differs significantly from natural conditions. This divergence is also reflected in the composition of the bacterial community. This may explain the observed similarity in microbiota composition across organs and the absence of organ-specific effects in the AQ_J site, a pattern similar to those observed in invaded regions. It is also noteworthy that lionfish invasions in the Atlantic Ocean and Caribbean Sea are largely attributed to releases from aquariums. Such conditions may have shaped the microbiota of released lionfish, resulting in differences from their wild counterparts. While data for direct comparisons are unavailable, this highlights the potential role of aquarium environments in influencing microbiota composition in ways that could facilitate invasion success. Moreover, the environment serves as a microbial reservoir to which organisms are continuously exposed, facilitating the acquisition and exchange of bacteria between the host and its surroundings. Stevens and Olson (2013) demonstrated that eggs extracted from the ovaries of pregnant LF were free of bacteria, indicating the absence of vertical transmission and highlighting the environment’s role as a source of bacterial acquisition for the microbiota composition. The Sex factor was also found to significantly influence the microbiota of LF as reported previously in other marine fish. Most studies have shown sex-based differences in the microbiota of the intestinal organs (Montalvo & Vázquez, 2018; He & Zhang, 2019; Hernández & Arrieta, 2020). However, microbiota differences have also been documented in other organs, such as the gills (Wang & Liu, 2021), skin (González-Fernández & Navas, 2020) and liver (Zhao & Xu, 2022). These findings, similarly observed in LF, highlight the role of physiological and hormonal differences between sexes in shaping microbial communities, potentially influencing immune responses and adaptation to environmental conditions (Liu & Yu, 2020; Givens & Ransom, 2018).

### Lessepsian invasive fish (MS - RS)

Early stages of establishment of the lionfish (*Pterois miles*) in the Mediterranean was reported around 2012 from Lebanon (Bariche et al., 2013), although a single observation from Israel dates back to 1991 (Golani & Sonin, 1992). The second sighting off the Israeli coast occurred in 2013 within the northern Achziv marine reserve, and during 2015 the lionfish has progressed southward to central Israel (Stern & Rothman, 2018). By 2024, *P. miles* had spread westward to Rhodes and Sicily, successfully establishing populations throughout the eastern Mediterranean (Bottacini et al., 2024). Genetic studies have confirmed that this invasion originated from the Red Sea via the Suez Canal (Bariche et al., 2017; Dimitriou et al., 2019). The Suez Canal serves as one of the world’s busiest marine trade routes and provides easy access for non-indigenous species to migrate from the RS to MS through ballast water or as fouling communities on ships (Williams et al., 1988; Galil et al., 2015; Lavoie et al., 1999; Costello et al., 2022). The establishment of lionfish in the Mediterranean follows a well-documented pattern of biological invasion, where populations stabilize in nearby regions before expanding to more distant areas. This pattern is also characteristic of other invasive species, such as *Diadema setosum,* which have similarly followed the Lessepsian migration route through the Suez Canal (Zirler et al., 2023). During the establishment process, invasive populations grow, adapt to local environmental conditions, and increase their potential for further expansion (Kolar & Lodge, 2000; Zirler et al., 2023).

We monitored the presence and distribution of LF (*P. miles*) along the Israeli coast beginning in 2017, focusing on two sites, two seasons and three depth zones. Over the five years, we observed, in Israeli coast, a spreading process, north to south and from deeper to shallower waters (Fig 6A, B), similar to the findings of Stern & Rothman (2018). The initial invasion of LF along the northern coast of Israel (Achziv), can be attributed by the cyclonic circulation pattern characterizing the Levant Basin in the eastern Mediterranean (Pinardi et al., 2006); however, it is more likely explained by the continuous complex rocky habitats, which may serve as a suitable recruitment area and are characteristic of the initial habitats where LF populations first established in Cyprus, Turkey, and Greece (Stern & Rothman 2018). Notable differences in LF density and distribution were observed between seasons, with higher numbers recorded in the fall. These differences may be driven by adaptations to environmental changes, such as temperature, or by biological factors like predation, food supply, and reproductive cycles. A similar pattern was previously found for other invasive fish which are more abundant in rocky reefs during the fall, when water temperature is higher (Lazarus et al., 2022). These dynamics can lead to shifts in habitat preferences, like increased depth or more sheltered areas (Corey, 2016). Seasonal health status may also play a role in fish distribution and behavior; for example, Minnow fish species exhibited significantly better health status (lower HAI score) and lower parasite abundance and diversity in autumn compared to spring (Almeida et al., 2024). The seasonal trends observed in LF densities and distribution indicate a need for further research to better understand how environmental and biological factors affect their seasonal behavior and habitat preferences. The initial observations of the LF survey were recorded at a depth of around 45 meters before they began appearing in shallower waters (25 and 10m), similar to observations previously reported by Stern & Rothman (2018). This distribution pattern may suggest that LF initially established themselves in deeper which likely offered a more protected environment with reduced competition, abundant food resources, and lower predation pressure (jones et al., 2014; Tomáš et al.,2018, Chaikin et al., 2022). Airey et al., (2023) demonstrated the connection between deep and shallow habitats by examining the post-settlement dispersal patterns of invasive LF. The study found that approximately 34.5% of the LF population in shallow waters consisted of individuals who had migrated from deeper habitats. In comparison, only about 4% had moved in the opposite direction, from shallow to deep waters. This pattern suggests that the primary direction of LF dispersal after settlement is from deeper regions to shallower habitats, emphasizing the role of deep-water areas as potential initial settlement zones before LF spreads into more coastal regions. In addition, their ability to exploit a range of depths likely contributes to their success as an invasive species (Kletou et al., 2016).

An increasing trend in the average body size of LF has been observed over the years in surveys conducted in the Mediterranean Sea. The average length (TL) of LF sampled in the RS region (where LF are native) and in CA (an earlier invasion area compared to MS) was approximately 30 cm. In contrast, the average length of LF sampled in the MS region was significantly smaller and only after five years (in 2022) did the LF population in this region reach this size. This observation may indicate the species’ establishment process and thriving in newly invaded regions. Furthermore, studies show that larger body sizes offer several advantages, including improved resources competition, enhanced survival in various environments, and increased reproductive rates. As a result, species with larger body sizes are generally more successful as invaders. (Roy et al.,2002; Schröder et al., 2009).

Dietary flexibility plays a crucial role in the success of invasive species as it enables them to exploit new resources in diverse environments. Previously studies that compared Lessepsian carnivorous invaders and their counterparts in the RS showed significant differences in isotopic patterns. These differences were more pronounced in *S. rubrum*, which is an earlier invader, compared to *Pterois miles* (Tsadok et al., 2023). A similar trend was seen in the calculated TP, where significant differences appeared only in the earlier-established invaders (Fig. 7c). In contrast, no difference was seen in TP of LF, which may suggest that the LF are still in the process of establishment and adaptation into the new environment. However, the TP of Caribbean LF was significantly lower from both MS and RS, which again emphasizes the environmental factor. In both *S. rubrum* (carnivores) and *s. rivulatus*, (herbivorous) the TP was higher in the RS than in the MS. This reflects how the Mediterranean environment influences food web dynamics and necessitates dietary adjustments by invasive fish during their acclimatization and suggests that Lessepsian species capable of dietary adaptation may be more likely to become successful colonizers (Golani, 1993). It is important to note that fish size was not considered in the TP analyses in this study and it may have affected our results, since fish diet, and particularly LF’s diet shifts as they grow (Dahl & Patterson 2014). Continued study of the LF in the Mediterranean Sea will be able to track the shifts that appear over time in isotope patterns and TP.

#### Photobacterium

*Photobacterium* is widely found in marine environments, where it plays a significant role in ecosystem dynamics, particularly through its interactions with marine fish. This genus, part of the *Vibrionaceae* family, is notable for its symbiotic and pathogenic interactions with various marine organisms (see review: Labella et al., 2017). Symbiotically, certain *Photobacterium* species, such as *P. phosphoreum* and *P. leiognathi*, produce bioluminescence in light organs of fish, contributing to communication, predation, camouflage, and predator protection. Additionally, these species contribute to nutrient cycling by releasing essential fatty acids, which provide nutritional resources within marine food webs and support broader ecological interactions (Moi 2017; Urbanczyk, 2011, Zarubin, 2012). However, much of the literature on *Photobacterium* species focuses on their potential pathogenicity. This genus is frequently linked to diseases in various marine species, especially fish. For instance, *P. damselae*, particularly its subspecies *P. damselae and P. piscicida*, poses significant threats to marine aquaculture and wild fish populations. These bacteria are responsible for causing photobacteriosis, a disease characterized by skin lesions, septicemia, and high mortality rates among affected fish, such as groupers and sea bream. (Rivas et al., 2013; Labella et al., 2017, Austin & Austin, 2016).

In our study, *Photobacterium* was found in all samples and constituted a dominant component of the LF microbiome core. However, compared to the RS region, the relative abundance of *Photobacterium* was significantly higher in the MS, where it exceeded 50% across all examined organs. This high abundance may explain the lower number of taxa (Genus level) observed in the MS compared to the RS region, in all examined organs (Fig 4). Interestingly, further comparative analysis of other Lessepsian invasive fish species, *S. rubrum S. rivulatus,* revealed a similar trend, with a significantly higher relative abundance of Photobacterium in the fish organs from the MS compared to the RS region. Invasive species may acquire beneficial microbes from co-occurring native species, which can accelerate their adaptation to local conditions and reduce invasion lag times (Martignoni & Kolodny, 2023).

Two dominant sequences were identified among the Photobacterium found in the MS and RS samples, corresponding to *Photobacterium damselae* subsp*. damselae*, according to BLAST results from NCBI. Interestingly, despite identifying the pathogenic species, no external or internal signs of disease were observed in the fish analyzed. This aligns with similar observations in additional studies on wild Mediterranean fish and sharks, where Photobacterium species presence did not correlate with visible disease symptoms (Meron et al., 2020; Bregman et al, 2023)). The high relative abundance of Photobacterium in Mediterranean lionfish (LF) raises further questions about its potential role in the fish’s resistance and adaptation. For instance, are lionfish resistant to these pathogens, or do these bacteria contribute to their acclimatization to new environmental conditions, thereby enhancing their success as invasive species? These questions underscore the need for further research to explore the dynamics and ecological mechanisms underlying the interactions between lionfish and Photobacterium.

### Summary

This study focuses on the microbiota of the lionfish (LF), characterizing its species-specific core microbiome and identifying habitat-specific markers while discussing its potential role in invasion and establishment. The findings suggest that lionfish (LF) are still establishing themselves in the Mediterranean environment but are exhibiting a growing presence. The relatively recent invasion of LF from the Red Sea to the Mediterranean offers a unique opportunity to monitor microbial and ecological establishment processes, including population distribution, body size, and dietary patterns. This understanding can offer valuable insights into ecological impacts and support the development of effective management strategies for invasive species.

## Acknowledgements

We would like to thank the staff at Caribbean Netherlands Science Institute (CNSI), The Gottesman Family Israel Aquarium, Jerusalem and Morris Kahn Marine Research Station (MKMRS) for their assistance and support.

